# Protocol for rapid allelic discrimination qPCR genotyping of the *Winnie* mouse model

**DOI:** 10.64898/2026.02.17.704640

**Authors:** Behzad Mansoori, Chengyu Liang

**Affiliations:** Program in Molecular and Cellular Oncogenesis, The Wistar Institute, Philadelphia, PA, 19104, USA

## Abstract

Winnie mice are a widely used in vivo model of inflammatory bowel disease carrying a missense mutation in the Muc2 gene. Here, we present a protocol for genotyping Winnie mice using TaqMan allelic discrimination quantitative PCR. We describe tissue collection, rapid crude DNA extraction, probe-based amplification with dual-labeled fluorophores, and fluorescence-based genotype calling in a single reaction. This protocol enables qualitative SNP genotyping without post-amplification processing and can be readily adapted to other defined point mutations.

**Graphical abstract:** 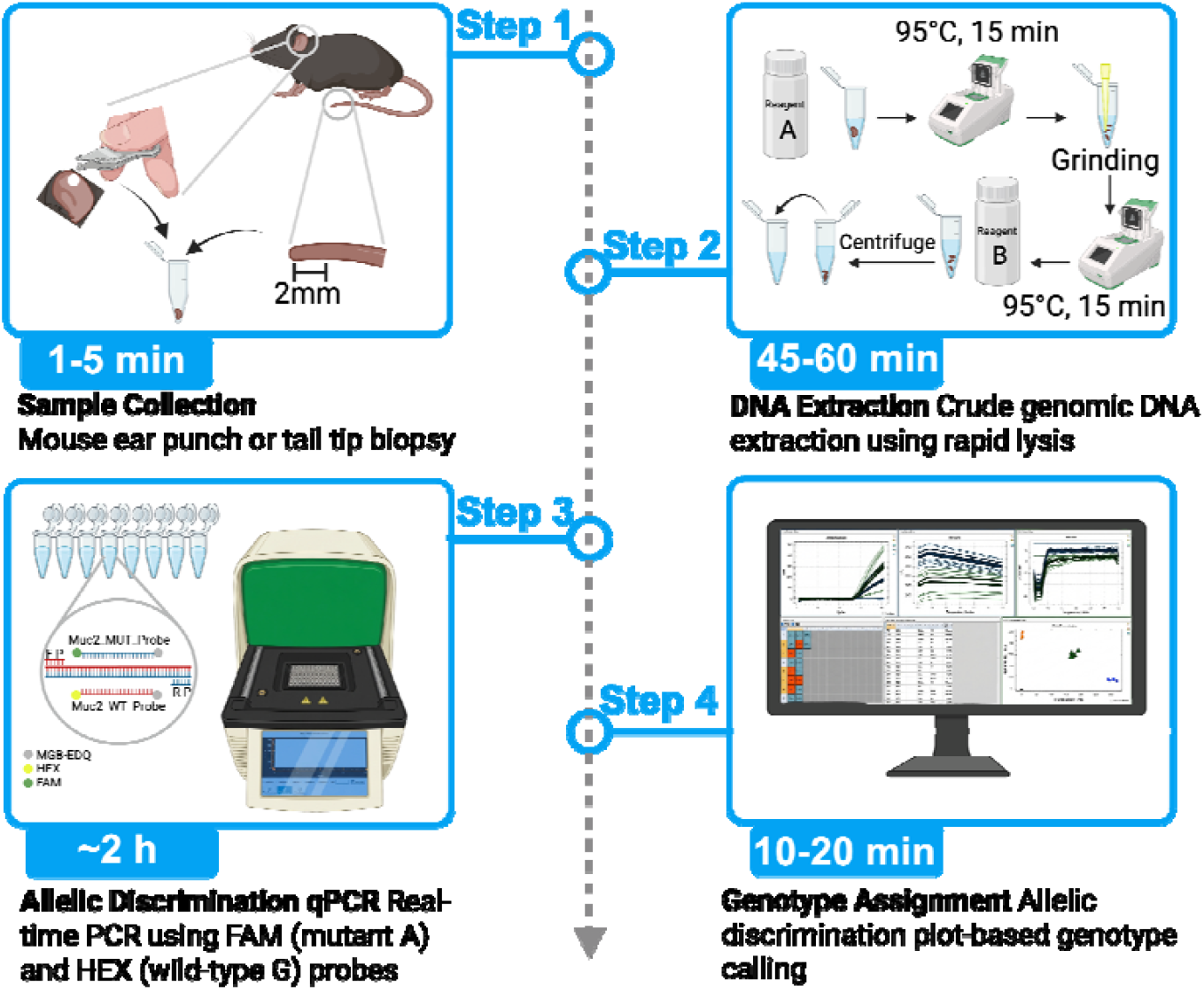

**Highlights:**

- Allelic discrimination qPCR protocol for genotyping the *Muc2* p.Cys52Tyr mutation using dual-labeled hydrolysis probes
- Enables rapid discrimination of wild-type, heterozygous, and mutant alleles in a single reaction
- Compatible with standard real-time PCR instruments and requires no post-PCR processing
- Supports high-throughput genotyping from crude DNA with minimal hands-on time

## Before you begin

Ulcerative colitis (UC) is a chronic inflammatory disorder characterized by mucosal injury, epithelial barrier dysfunction, and dysregulated immune responses^1^. Winnie mice harbor a G-to-A missense mutation (G9492A) in the *Muc2* gene that disrupts proper folding and assembly of the MUCIN-2 protein in the endoplasmic reticulum, resulting in spontaneous, progressive colitis that is largely localized to the colon^2^. This model recapitulates key epithelial, immunological, and microbiota-associated features observed in UC patients and is widely used to study chronic intestinal inflammation^2–5^. Compared with chemically induced models such as dextran sulfate sodium (DSS), which rely on acute epithelial injury, the Winnie model provides a genetically driven platform for investigating disease initiation, progression, and therapeutic responses over time^4^.

Accurate determination of *Muc2* genotype is essential for distinguishing homozygous mutant, heterozygous mutant, and wild-type (WT) animals, as disease severity, penetrance, and experimental outcomes are strongly influenced by gene dosage^2,4^. Routine colony management therefore requires a genotyping strategy that is reliable, rapid, and scalable, particularly when managing large breeding cohorts or time-sensitive experimental studies. The protocol described here was developed for routine genotyping of Winnie mice using TaqMan allelic discrimination quantitative PCR (qPCR)^6,7^. In our studies, this approach was applied to ear punch and tail tip biopsies collected from breeding and experimental cohorts. The assay employs two allele-specific hydrolysis probes labeled with FAM (mutant A allele) and HEX (WT G allele), allowing discrimination of all three genotypes within a single qPCR reaction without post-amplification processing. Crude genomic DNA prepared using a rapid lysis-based extraction method is sufficient for reliable genotype assignment.

**Note:** standard curves are not required for TaqMan allelic discrimination qPCR assays. This approach is designed for qualitative SNP genotyping rather than quantitative DNA measurement. Genotypes are assigned based on allele-specific fluorescence clustering (e.g., FAM for mutant and HEX for WT) rather than amplification efficiency or copy number. Accordingly, this workflow does not rely on standard dilutions or curve fitting commonly used in gene expression or microbial load assays.

Before beginning the protocol, ensure that allele-specific primers and probes have been ordered, a real-time PCR instrument capable of detecting FAM and HEX fluorescence channels is available, and reagents for rapid DNA extraction are prepared as described below. Although optimized for genotyping the *Muc2* allele, this allelic discrimination approach can be adapted to other defined single-nucleotide polymorphisms (SNPs) with appropriate probe design.

### Innovation

Accurate genotyping the Winnie mouse strain is essential for studies of chronic colitis, as disease severity and experimental outcomes are strongly influenced by *Muc2* gene dosage. Traditional genotyping approaches typically involve PCR amplification followed by Sanger sequencing or gel-based assays. Although reliable, these workflows are time-intensive, require post-PCR processing, and are not well suited for routine or large-scale colony management.

To address these limitations, we developed an allelic discrimination qPCR workflow specifically configured for detection of the *Muc2* G-to-A point mutation in Winnie mice. This protocol integrates rapid lysis-based DNA extraction with dual-labeled, allele-specific hydrolysis probes to distinguish wild-type, heterozygous, and homozygous mutant alleles within a single closed-tube reaction. Genotype assignment is achieved directly from fluorescence signals without sequencing or gel electrophoresis. A key advancement of this protocol is its compatibility with crude DNA prepared from ear punch or tail tip biopsies and its implementation on standard real-time PCR platforms commonly available in research laboratories. The protocol reduces hands-on time and supports parallel processing of large sample numbers while maintaining clear genotype discrimination. With appropriate probe design, this approach can be adapted to other defined single-nucleotide variants, making it a practical solution for routine colony genotyping and related applications.

### Institutional permissions (if applicable)

All mouse tissues used in this study were obtained from animals maintained and handled in accordance with institutional and national guidelines for animal care and use. Animal procedures were performed in accordance with approval from the Institutional Animal Care and Use Committee (IACUC) at The Wistar Institute. Readers are responsible for obtaining all required institutional and regulatory approvals before performing experiments involving animals or animal-derived samples.

### Preparation steps

#### Preparing and storing primers and probes

##### Timing: Typically 1-2 working days (may vary by supplier and location)

This section describes how to order, resuspend, aliquot, and store primers and allele-specific hydrolysis probes for TaqMan allelic discrimination qPCR. Proper handling and storage of probes are critical to maintain fluorescence performance and genotype discrimination.

1. Order the forward and reverse primers and the allele-specific TaqMan hydrolysis probes listed in **Table 1** from a commercial oligonucleotide supplier. **Note:** Probes bind the SNP site with allele specificity. The HEX-labeled probe detects the WT *Muc2* allele (G), and the FAM-labeled probe detects the mutant allele (A). Probes contain a 3’ NFQ-MGB to minimize background fluorescence.
2. Upon receipt, briefly centrifuge primer and probe tubes to collect material at the bottom.
3. Resuspend primers and probes in nuclease-free water according to the manufacturer’s instructions to generate concentrated stock solutions (e.g., 100 µM).
4. Mix thoroughly by gentle pipetting and incubate at room temperature (RT) for 5 min to ensure complete dissolution.
5. Prepare single-use or small-volume aliquots of primers and probes to minimize repeated freeze-thaw cycles.
6. Store primer stock solutions at −20°C.
7. Store probe aliquots at −20°C protected from light.

**Table 1.**
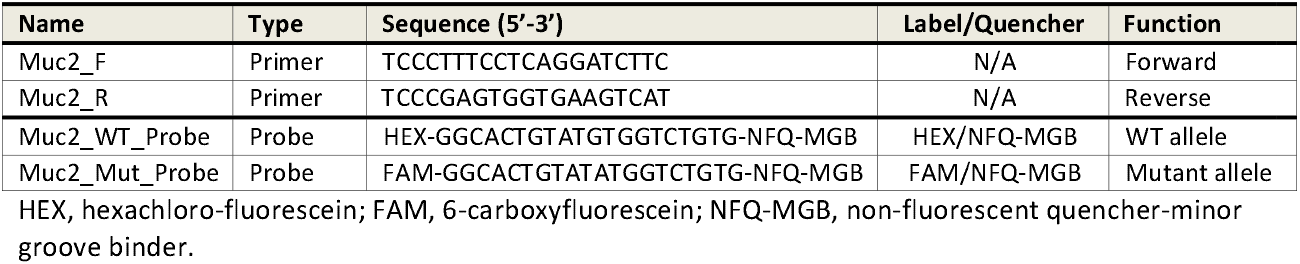
Primers and probes for TaqMan SNP genotyping of Winnie mouse mutation.

##### Storage conditions

| Reagent | Storage temperature | Maximum storage time |
| --- | --- | --- |
| Primers | -20°C | ≥ 12 months |
| Hydrolysis probes (FAM/HEX-labeled) | -20°C, protected from light | ≥ 12 months |

**Note:** Hydrolysis probes are light-sensitive. Repeated freeze-thaw cycles or prolonged exposure to light can reduce fluorescence intensity and compromise allelic discrimination.

#### Preparing mouse tissue samples for genotyping

##### Timing: 1-5 min per sample (may vary depending on sample number)

This section describes preparation and handling of mouse tissue samples for downstream allelic discrimination qPCR genotyping. Ear punch or tail tip biopsies can be collected in advance and stored prior to DNA extraction.

8. Collect mouse tissue samples using one of the following approaches based on animal age and institutional guidelines.
  a. Ear punch samples: at weaning age or older, collect a single ear punch biopsy using a sterile ear punch tool.
  b. Tail tip samples: from pre-weaning mice, collect a 1.5-2 mm distal tip using sterile scissors.
9. Place each tissue sample immediately into a labeled PCR tube or microcentrifuge tube designated for DNA extractions. **Critical:** Use sterile instruments for each animal, or disinfect tools between animals, to prevent cross-contamination.
10. If DNA extraction will be performed immediately, proceed directly to the extraction steps described below.
11. If extraction will be delayed, store tissue samples at −20°C until use.

**Note:** Ear punch and tail tip samples yield sufficient genomic DNA for TaqMan allelic discrimination qPCR without additional purification steps.

**Note:** Avoid repeated freeze-thaw cycles of tissue samples, as this may reduce DNA quality and affect downstream amplification.

#### Preparing lysis buffer for crude DNA extraction

##### Timing: 10-15 min

This section describes preparation of the alkaline lysis and neutralization buffers used for rapid extraction of genomic DNA from mouse ear punch or tail tip biopsies. These buffers enable efficient DNA release without column purification and are compatible with downstream TaqMan qPCR.

12. Prepare alkaline lysis buffer (Solution A) according to the manufacturer’s instructions (PreciGene™ Quick Extraction Kit) or equivalent alkaline lysis formulation.
13. Prepare neutralization buffer (Solution B) according to the manufacturer’s instructions.

##### Storage conditions

| Solution | Storage temperature | Maximum storage time |
| --- | --- | --- |
| Alkaline lysis buffer (Solution A) | Room temperature | As specified by manufacturer |
| Neutralization buffer (Solution B) | Room temperature | As specified by manufacturer |

**Note:** If preparing homemade alkaline lysis buffer, ensure compatibility with downstream qPCR and validate performance using known genotype controls before routine use.

## Key resources table

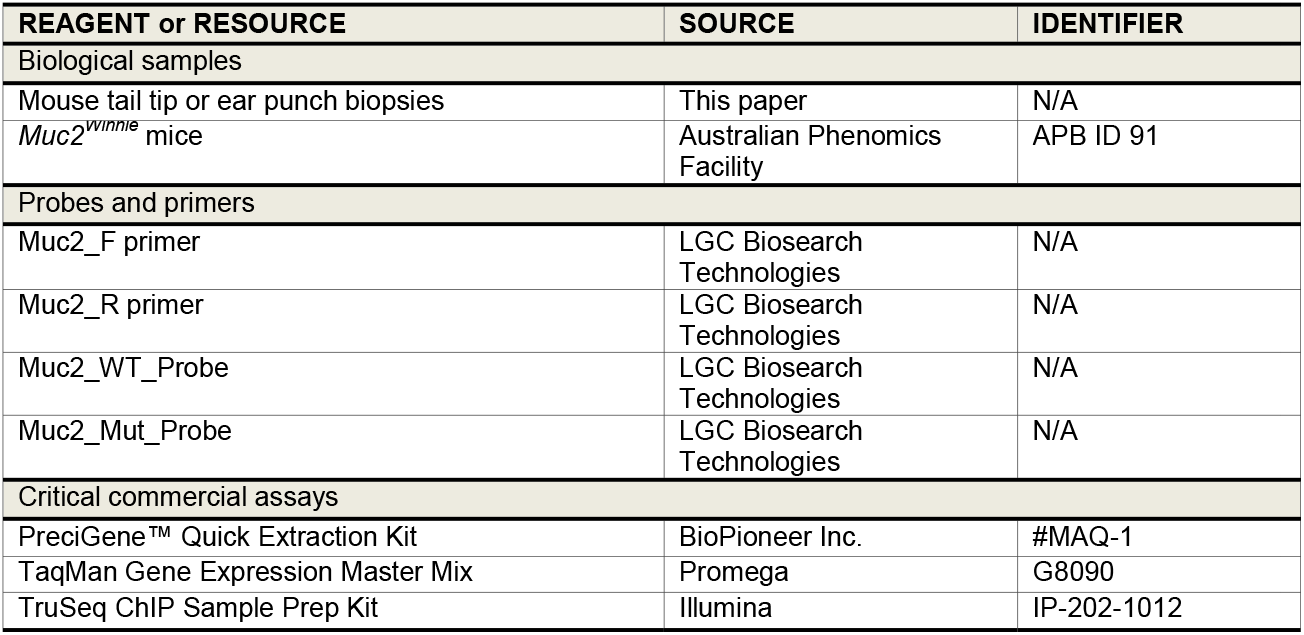
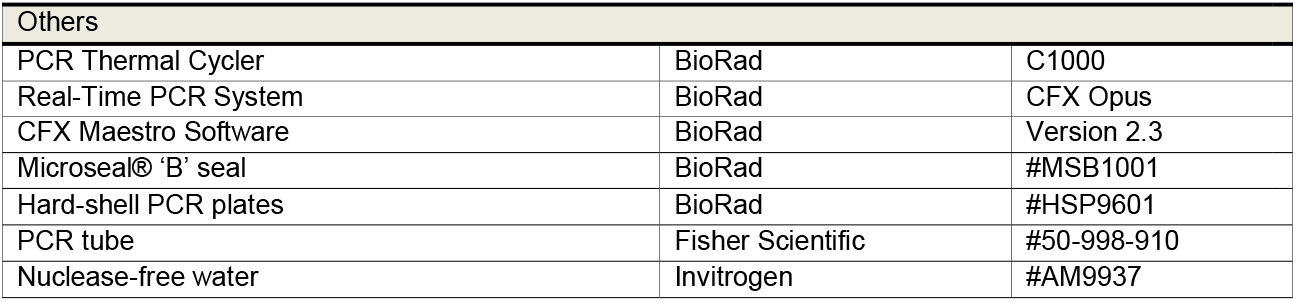

| REAGENT or RESOURCE | SOURCE | IDENTIFIER |
| --- | --- | --- |
| Biological samples |  |  |
| Mouse tail tip or ear punch biopsies | This paper | N/A |
| *Muc2*<sup>winnie</sup> mice | Australian Phenomics Facility | APB ID 91 |
| Probes and primers |  |  |
| Muc2_F primer | LGC Biosearch Technologies | N/A |
| Muc2_R primer | LGC Biosearch Technologies | N/A |
| Muc2_WT_Probe | LGC Biosearch Technologies | N/A |
| Muc2_Mut_Probe | LGC Biosearch Technologies | N/A |
| Critical commercial assays |  |  |
| PreciGene™ Quick Extraction Kit | BioPioneer Inc. | #MAQ-1 |
| TaqMan Gene Expression Master Mix | Promega | G8090 |
| TruSeq ChIP Sample Prep Kit | Illumina | IP-202-1012 |

| Others |  |  |
| --- | --- | --- |
| PCR Thermal Cycler | BioRad | C1000 |
| Real-Time PCR System | BioRad | CFX Opus |
| CFX Maestro Software | BioRad | Version 2.3 |
| Microseal® 'B' seal | BioRad | #MSB1001 |
| Hard-shell PCR plates | BioRad | #HSP9601 |
| PCR tube | Fisher Scientific | #50-998-910 |
| Nuclease-free water | Invitrogen | #AM9937 |

## Step-by-step method details

### Extract genomic DNA from tissue samples

#### Timing: 45-60 min (may vary depending on sample number and operator experience)

This section describes a rapid, lysis-based procedure for extracting genomic DNA from mouse ear punch or tail tip biopsies for downstream TaqMan allelic discrimination qPCR. The method yields DNA of sufficient quality for SNP genotyping without column purification or precipitation steps and is optimized for high-throughput processing.

**Critical:** Use sterile instruments and clearly labeled tubes throughout to prevent cross-contamination between samples.

1. Add alkaline lysis buffer to tissue samples.
  a. Place each ear punch or tail tip biopsy in a labeled PCR tube.
  b. Add 30 µl of alkaline lysis buffer (Solution A, PreciGene™ Quick Extraction Kit) directly to each tube.
  c. Incubate samples at 95°C for 15 min using a thermal cycler or heat block.
2. Mechanically dissociate tissue.
  a. Briefly centrifuge tubes to collect condensation.
  b. Using a sterile pipette tip, grind the tissue with a twisting motion 3-5 times to aid dissociation. **Note:** Complete mechanical disruption improves DNA release and downstream amplification consistency.
3. Complete lysis and neutralize the reaction.
  a. Return samples to 95°C for an additional 15 min.
  b. Cool samples to room temperature for 2 min and briefly centrifuge.
  c. Add 30 µl of neutralization buffer (Solution B) and mix thoroughly by pipetting.
4. Clarify lysates and prepare DNA for qPCR
  a. Centrifuge samples at 13,000 rpm for 1 min to pellet debris.
  b. Transfer 10 µl of the clear supernatant to a fresh PCR tube.
  c. Dilute the DNA 1:10 with nuclease-free water.
  d. Use diluted DNA immediately for qPCR or store at −20°C.

**Pause point:** Store diluted DNA at −20°C for up to several weeks.

**Troubleshooting:** If amplification fails or clustering is ambiguous, see Troubleshooting Problems 1-2.

### TaqMan allelic discrimination qPCR

#### Timing: 20~30 min (plate setup) + ~ 2 h (run)

This section describes setup and execution of TaqMan allelic discrimination qPCR for genotyping the *Muc2* G-to-A SNP using dual-labeled hydrolysis probes. Allele-specific fluorescence signals generated during amplification enable genotype assignment (**Figure 1**).

**Figure 1:**
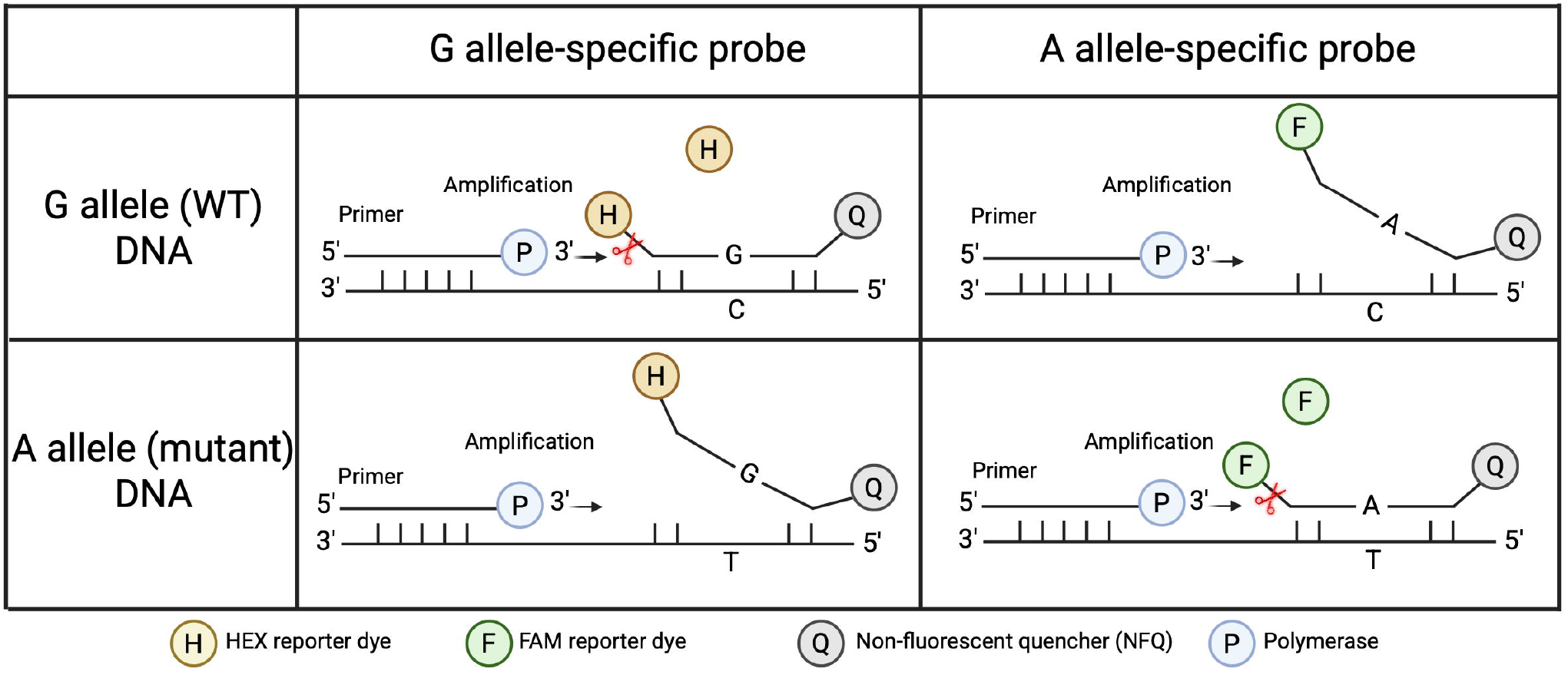
Schematic of TaqMan allelic discrimination for detection of the *Muc2* G-to-A mutation. The assay uses two allele-specific hydrolysis probes in a single-qPCR reaction to distinguish *Muc2* genotypes. A HEX-labeled probe detects the WT *Muc2* G allele, whereas a FAM-labeled probe detects the mutant A allele. During amplification, the 5’-nuclease activity of the DNA polymerase cleaves a perfectly matched probe, separating the reporter dye from the quencher and generating fluorescence. In WT samples, only the HEX-labeled probe is cleaved, whereas in mutant samples, only the FAM-labeled probe generates signal. Probes that are mismatched to the target allele remain uncleaved and do not produce fluorescence.

**Table 2.** qPCR reaction master mix per sample.

| Reagent | Volume (μl) | Final concentration |
| --- | --- | --- |
| TaqMan Universal PCR Master Mix (2x) | 5.0 | 1x |
| Muc2_F primer | 1.0 | 1 μM |
| Muc2_R primer | 1.0 | 1 μM |
| Muc2_WT_Probe (HEX) | 0.5 | 0.20 μM |
| Muc2_Mut_Probe (FAM) | 0.5 | 0.20 μM |
| DNA (1:10 dilution) | 1.0 | – |
| Nuclease-free water | 1.0 | – |
| Total | 10.0 | – |

5. Prepare the qPCR master mix (Table 2)
  a. Thaw the 2x TaqMan Universal PCR Master Mix, primers, and probes on ice.
  b. Mix reagents gently by pipetting or brief vortexing and centrifuge briefly.
  c. Prepare a master mix sufficient for all samples plus at least 10% excess.
  d. Dispense reagents according to Table 2 to achieve a final reaction volume of 10 µl per well. **Critical:** Protect probes from light during setup. **Critical:** Include no-template controls and known genotype controls (homozygous WT, heterozygous, homozygous mutant) in every run. C57BL/6 mice can be used as a homozygous WT reference. Previously sequenced samples with confirmed genotypes should be used as positive controls for mutant and heterozygous alleles.
6. Dispense reactions into a PCR plate.
  a. Aliquot 9 µl of master mix into each well.
  b. Add 1 µl of 1:10 diluted genomic DNA to each sample well.
  c. Seal the plate and briefly centrifuge to remove bubbles and collect liquid at the bottom of wells.
7. Configure the qPCR run in the instrument software.
  a. Select fluorescence detection for both FAM and HEX channels.
  b. Disable unused fluorophores (e.g., ROX, Cy5).
  c. Assign the SNP target name (i.e., *Muc2*) and remove unrelated default targets.
  d. Confirm that at least one well contains a valid sample with both fluorophores enabled.
  e. Set the qPCR run mode to a 2-step amplification protocol. **Note:** Configuration steps may vary slightly depending on instrument software (e.g., Bio-Rad CFX Manager or equivalent). **Critical:** Incorrect channel selection can prevent accurate genotype calling.
8. Run the qPCR thermal cycling program (Table 3) **Note:** Acquire fluorescence for both FAM and HEX channels during the 60°C annealing/extension step. **Troubleshooting:** If fluorescence signals are weak or absent, see Troubleshooting Problems 1 and 5.

**Table 3.** Thermal cycling conditions.

| Step | Temperature | Time | Cycles |
| --- | --- | --- | --- |
| Initial denaturation | 95°C | 10 min | 1 |
| Denaturation | 95°C | 15 s | 40 |
| Annealing/Extension | 60°C | 60 s |  |

**Note:** Acquire fluorescence for both FAM and HEX channels during the 60°C annealing/extension step.

### Review amplification curves and assess run quality

#### Timing: 5-10 min

9. Review amplification curves after completion of the qPCR run.
  a. Display amplification plots in linear scale view.
  b. Confirm that positive samples show a sigmoidal (exponential) amplification curve (**Figure 2**).
  c. Consider samples with Ct < 35 and a clear sigmoidal curve as successfully amplified.
  d. Treat samples with Ct > 35 or non-sigmoidal curve shapes (e.g., linear drift or background noise) as undetermined and exclude them from genotype calling.
  **Note:** Baseline-subtracted fluorescence values should be used when assessing curve shape consistency.

**Figure 2.**
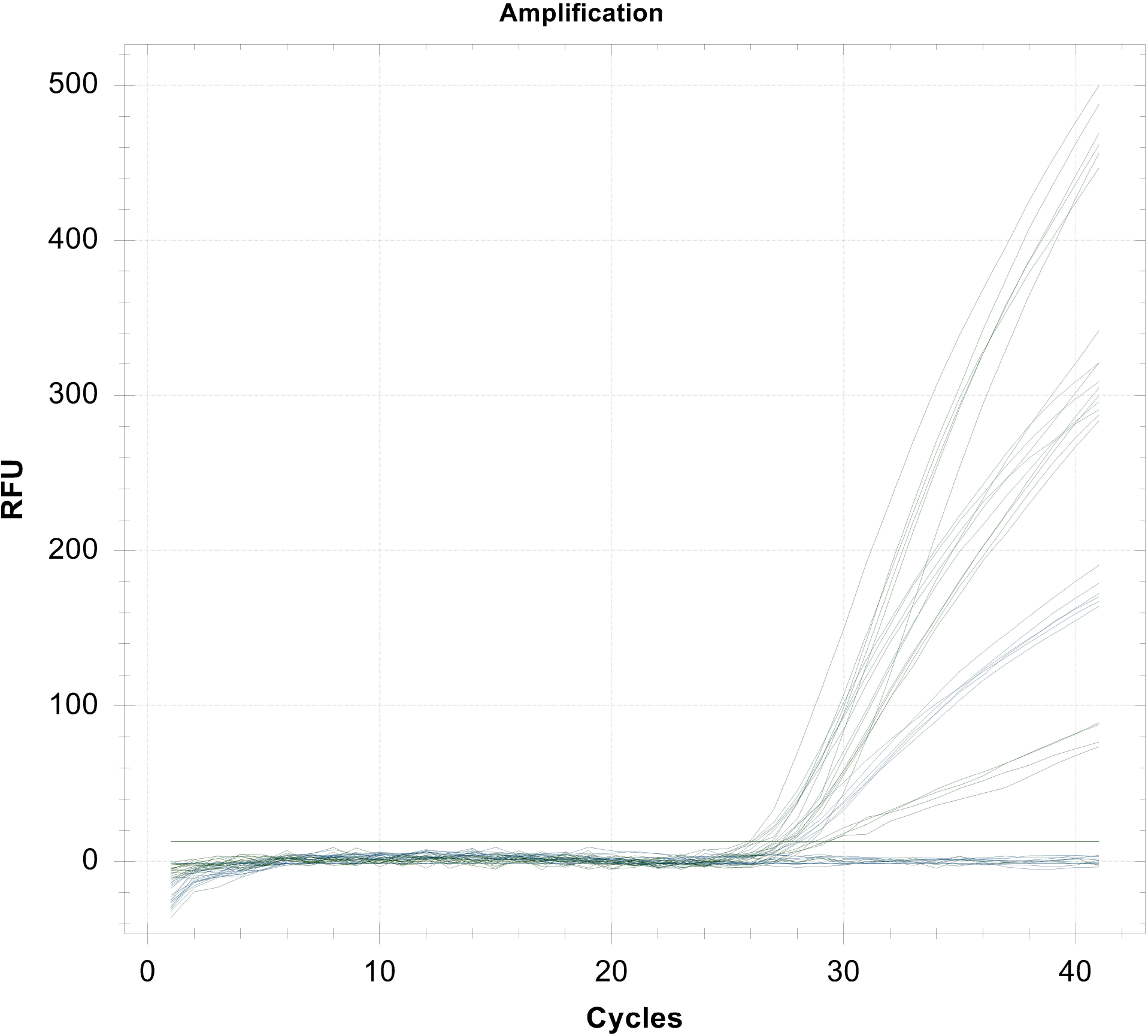
Representative amplification curves from allelic discrimination qPCR. Real-time qPCR amplification curves displayed in linear scale, showing fluorescence intensity (RFU) plotted against cycle number. Samples with successful target amplification exhibit characteristic sigmoidal fluorescence trajectories and cross the threshold before 35 cycles, whereas no-template controls and samples with late or irregular signals show minimal or inconsistent fluorescence.

### Expected outcomes

When this protocol is performed successfully, real-time qPCR amplification curves generated from mouse genomic DNA display characteristic sigmoidal fluorescence trajectories when visualized in linear scale (Figure 2). Samples containing amplifiable *Muc2* DNA typically show exponential signal increases that cross the threshold before 35 cycles, whereas no-template controls exhibit minimal background fluorescence. Samples with late or irregular signals beyond this range may not yield reliable genotype assignments.

Allelic discrimination analysis produces clear separation of samples based on allele-specific fluorescence intensity (Figure 3). Under standard conditions, samples cluster into four groups corresponding to homozygous WT (*Muc2* G/G; HEX signal only), homozygous mutant (*Muc2* A/A; FAM signal only), heterozygous (*Muc2* G/A; combined FAM and HEX signals), and no-template controls (low signal in both channels) (Table 4). These clusters are readily visualized when allelic discrimination plots are displayed using Cartesian coordinates.

**Table 4.**
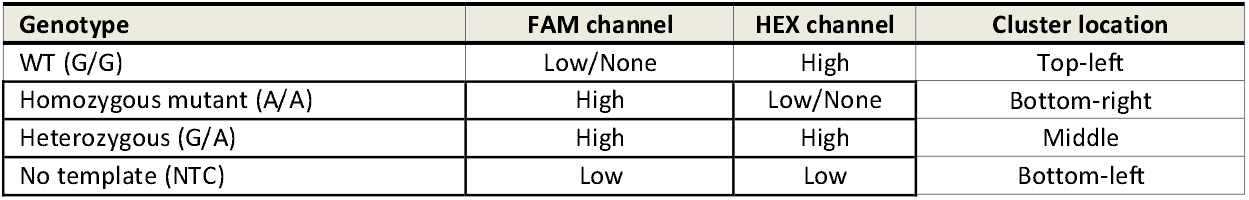
Expected allelic discrimination readout.

**Figure 3.**
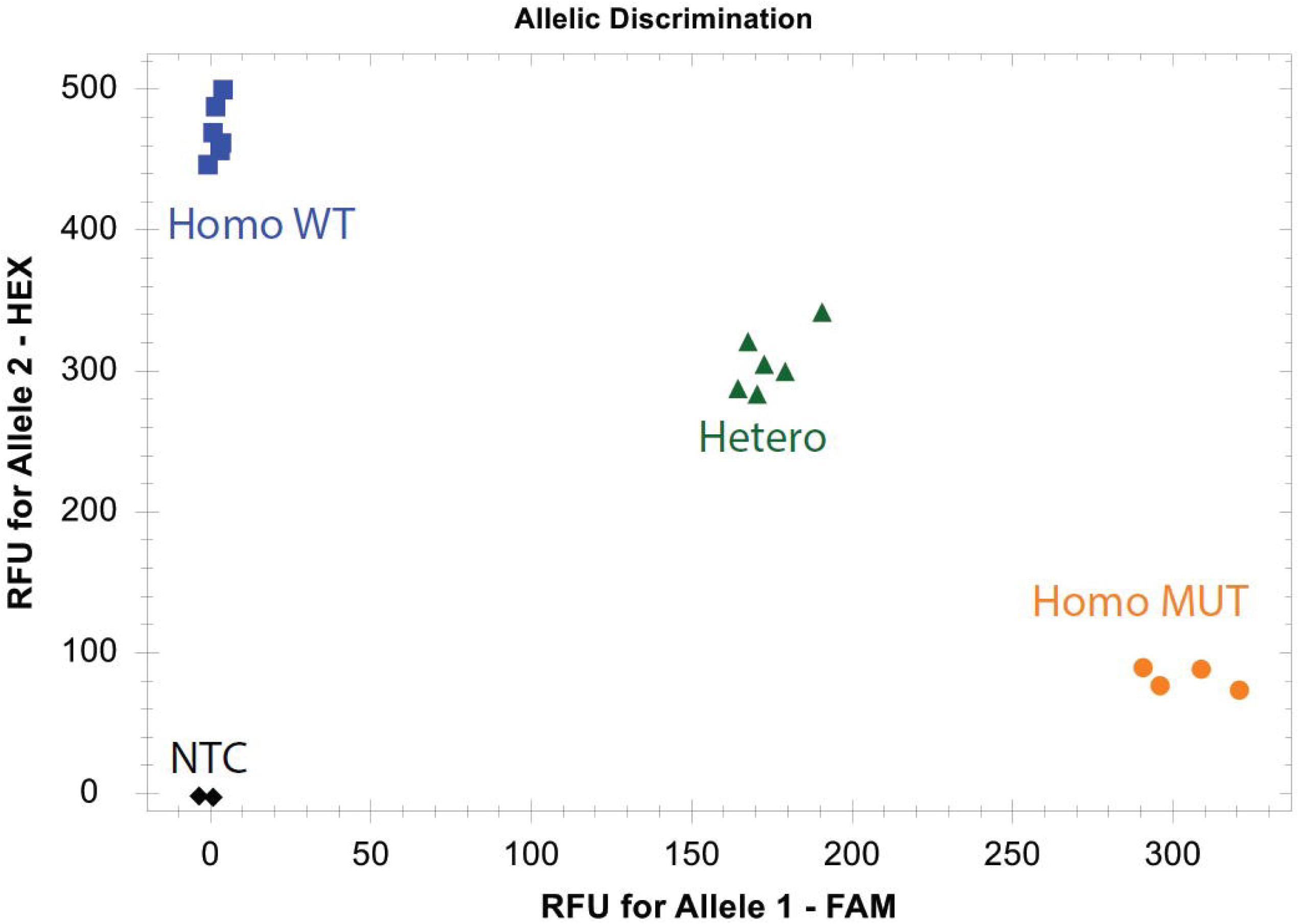
Representative allelic discrimination plot for *Muc2* genotyping. Scatter plot showing allele-specific fluorescence intensity displayed in Cartesian coordinates, with FAM signal on the x-axis and HEX signal on the y-axis. Samples segregate into distinct clusters corresponding to homozygous mutant (Homo Mut), homozygous WT (Homo WT), heterozygous, and no-template control (NTC) groups. Symbols and colors indicate genotype categories as shown.

Using crude DNA extracted from ear punch or tail tip biopsies, genotype clusters are typically well resolved without evidence of cross-signal between allele-specific probes. Genotype assignments are consistent with independent sequencing-based genotyping when tested on reference samples. The protocol therefore produces reproducible qualitative genotyping data suitable for routine use in colony management and experimental cohort assignment.

### Limitations

This protocol is designed for rapid genotyping of a defined single-nucleotide variant and has several limitations that should be considered. First, TaqMan allelic discrimination qPCR is inherently targeted and can only detect the predefined *Muc2* G-to-A mutation encompassed by the probe-binding region. It cannot identify additional or unexpected sequence variants, insertions, deletions, or structural alterations. When genetic drift, spontaneous mutations, or background strain variability are concerns, confirmatory sequencing remains necessary. Second, genotype assignment depends on the quality of the input DNA and the clarity of fluorescence signal clustering. Crude DNA extracts that are highly degraded, contain inhibitory contaminants, or yield borderline amplification (e.g., Ct values near or above 35) may produce ambiguous clustering. Such samples may require repeat extraction, rerunning the assay, or exclusion from analysis. Third, allelic discrimination relies on stable fluorophore detection and consistent instrument performance. Differences in real-time PCR platforms, optical calibration, fluorophore sensitivity, or plate sealing can influence fluorescence intensity and cluster separation. Maintaining consistent instrumentation, reaction volumes, and run conditions is therefore important for reproducibility across experiments. Finally, accurate genotyping assumes careful tissue collection and sample handling. Cross-contamination during ear punch or tail tip sampling, mislabeling, or plate setup errors can lead to incorrect genotype calls. In addition, this protocol is limited to biallelic discrimination at a single locus and is not suitable for multiplex genotyping or high-resolution mapping of multiple variants in parallel.

### Troubleshooting

#### Problem 1

No amplification or absence of fluorescence signal

Related to: Step 9 (Review amplification curves and assess run quality); Step 8 (qPCR thermal cycling).

#### Potential solution

- Confirm that both allele-specific probes (FAM- and HEX-labeled) and primers were included at the correct concentrations during master mix preparation.
- Verify that the thermal cycling program and fluorescence acquisition settings are correctly configured for FAM and HEX detection.
- Assess DNA quality and input amount. Highly degraded DNA or very low template input may prevent amplification.
- If inhibitors from crude DNA extraction are suspected, dilute the DNA sample (e.g., 1:10) and repeat the reaction.

#### Problem 2

High background fluorescence or nonspecific amplification curves

Related to: Step 9 (Review amplification curves)

#### Potential solution

- Inspect amplification curves for nonspecific signal or drift rather than sigmoidal amplification.
- Reduce template input if excessive DNA is used, as high DNA concentrations may increase background signal.
- Use freshly prepared master mix and ensure thorough mixing of reaction components.
- If nonspecific amplification persists, primers may require redesign to improve specificity.

#### Problem 3

Fluorescence signal detected in no-template control (NTC).

Related to: Step 8 (qPCR setup and run)

#### Potential solution

- Treat this result as evidence of contamination and discard the run.
- Prepare fresh reagents and master mix using new filter tips in a clean workspace.
- Physically separate pre-PCR and post-PCR areas to minimize contamination risk.
- Always include at least one NTC per run to monitor contamination.

#### Problem 4

Ambiguous genotype clustering in allelic discrimination plots Related to: Step 10 (Allelic discrimination analysis)

Variability in fluorescence intensity or borderline amplification can lead to partial overlap between genotype clusters in the allelic discrimination plot. This is most commonly observed in samples with low-quality DNA, late amplification, or when appropriate genotype controls are not included in the run.

#### Potential solution

- DNA quality and input: re-extract DNA from the tissue sample or dilute the crude extract (e.g., 1: 10) to reduce inhibitory effects that can distort fluorescence signals.
- Ct proximity to cutoff: samples with Ct values near or above 35 may show unstable clustering. Re-running these samples with freshly prepared DNA can improve separation.
- Control inclusion: include at least one known WT, heterozygous, and mutant control per run to guide cluster assignment.
- Plot configuration: ensure that allelic discrimination plots are displayed in Cartesian coordinates and that only FAM and HEX channels are enabled.

#### Problem 5

Weak FAM or HEX signal despite successful amplification

Related to: Step 10 (Allelic discrimination analysis)

#### Potential solution

- Confirm that fluorophore detection channels are properly calibrated on the qPCR instrument.
- Minimize probe degradation by avoiding repeated freeze-thaw cycles and protecting probes from light exposure.
- Use freshly aliquoted probes stored at −20°C in the dark.
- Verify probe concentration and ensure uniform mixing before loading reactions.

### Resource availability

#### Lead contact

Further information and requests for resources and reagents should be directed to and will be fulfilled by the lead contact, Chengyu Liang.

#### Technical contact

Technical questions on executing this protocol should be directed to and will be answered by the technical contact, Behzad Mansoori.

#### Materials availability

This protocol does not generate new unique reagents.

#### Data and code availability

This protocol does not include datasets.

## Acknowledgments

The project was supported by The Wistar Institute Training Program (T32 CA009171) to B. Mansoori; NIH awards R01 CA140964, R01 CA262631 to C. Liang (PI). All graphic illustrations were created using BioRender.com under an institutional license.

## Author contributions

Conceptualization, B.M. and C.L.; methodology, B.M.; investigation, B.M.; supervision, C.L.; writing—original draft, B.M.; writing—review and editing, B.M. and C.L.

